# A physical neural mass model framework for the analysis of oscillatory generators from laminar electrophysiological recordings

**DOI:** 10.1101/2022.07.19.500618

**Authors:** Roser Sanchez-Todo, André M. Bastos, Edmundo Lopez Sola, Borja Mercadal, Emiliano Santarnecchi, Earl K. Miller, Gustavo Deco, Giulio Ruffini

## Abstract

Cortical function emerges from the interactions of multi-scale networks that may be studied at a high level using neural mass models (NMM), representing the mean activity of large numbers of neurons. In order to properly reproduce experimental data, these models require the addition of further elements. Here we provide a framework integrating conduction physics that can be used to simulate cortical electrophysiology measurements, particularly those obtained from multi-contact laminar electrodes. This is achieved by endowing NMMs with basic physical properties, such as the average laminar location of the apical and basal dendrites of pyramidal cell populations. We call this framework laminar NMM, or LaNMM for short. We then employ this framework to infer the location of oscillatory generators from laminar-resolved data collected from the prefrontal cortex in the macaque monkey. Based on the literature on columnar connectivity, we define a minimal neural mass model capable of generating amplitude and phase coupled slow (alpha/beta, 4–22 Hz) and fast (gamma, 30–250 Hz) oscillations. The synapse layer locations of the two pyramidal cell populations are treated as optimization parameters, together with two more LaNMM-specific parameters, to compare the models with the multi-contact recordings. We rank the candidate models using an optimization function that evaluates the match between the functional connectivity (FC) of the model and data, where the FC is defined by the covariance between bipolar voltage measurements at different cortical depths. The family of best solutions reproduces the FC of the observed electrophysiology while selecting locations of pyramidal cells and their synapses that result in the generation of fast activity at superficial layers and slow activity across most depths, in line with recent literature proposals.

**Highlights:**

- We provide a neural mass modeling formalism that includes a physical layer to simulate electrophysiology measurements.
- To analyze in-vivo data collected in the macaque monkey during a memory task, we propose a specific model with two coupled main circuits that can generate realistic electrophysiological signals in two important oscillatory regimes—the alpha/beta and the gamma bands.
- Physical elements in the model shed light on the generation of oscillations in the two regimes and on the relative power distribution of fast and slow oscillatory signals across cortical depth, which we show can be altered by the choice of the reference location or method.
- The model is contrasted with in-vivo data, with parameters adjusted by matching voltage statistics in the alpha/beta and gamma bands, leading to a solution with slow frequency components generated by synapses spanning most cortical layers and fast oscillations in superficial layers.
- The resulting formalism provides useful tools and concepts to analyze and model data, with implications for understanding altered oscillatory EEG activity in dementia, Alzheimer’s disease and other disorders with oscillatory features.

## 1. Introduction

Brain function results from interactions between specialized, spatially-distributed areas of brain networks [4]. For this reason, to explore the relationship between function and the underlying structure, the brain can be modeled as a complex and dynamic multi-scale network (for a review, see [3, 1]). Along these lines, several computational studies rely on large-scale neural mass models (NMM) networks to obtain insights into brain physiology and its underlying mechanisms [40, 29, 52, 8].

Jansen and Rit’s NMM [24] is an effective *lumped* mesoscale model of neuronal populations based on the work of Lopes Da Silva and van Rotterdam in the 1970s [33, 34, 51]. It describes cortical column dynamics by capturing relevant physiological features at the mesoscale, but in its original form, it has some limitations. For example, it can represent oscillations only in one specific frequency band for each parameter configuration [17], limiting its usefulness in modeling disorders with multifrequency alterations such as Alzheimer’s [49]. This can be remedied by adding more neuronal populations to the original model [55], as we do here. Furthermore, NMMs cannot *per se* generate measurements such as local field potentials (LFP) or derived quantities such as current source density (CSD) since these models are not mathematically embedded in a physical medium. They do, however, provide a handle on synaptic current sources and membrane potentials, where physics modeling can begin. While membrane potential may be sufficient for comparison with patch-clamp experiments, adding further physical structure is necessary to contrast model outputs with electrophysiological recordings such as LFPs, stereotactic EEG (SEEG), or, in whole brain network models, scalp electro- or magnetoencephalography (EEG or MEG).

The raw outputs of NMMs are the membrane potential alterations induced by each synapse and the consequent firing rates of each population in the model. They are determined by a set of parameters describing, e.g., the dynamics of post-synaptic potentials, the relationship between membrane potential alteration and firing rate, population connectivity, and external inputs. Several studies have employed rodent multi-unit activity (MUA), LFP, and CSD measurements to estimate or fit some of these parameters [56, 10, 31, 45]. The average membrane potential or firing rate of the pyramidal populations is typically used to compare model outputs with MUA [10], LFP [45], or CSD [31] measurements. Whole-brain computational studies use similar methods to simulate macroscopic electrophysiological recordings (e.g., EEG) in humans [40, 29, 46]. However, as discussed here, the connection between NMM membrane potential or firing rates with electrophysiology is not well defined. Unlike detailed neuron compartment models [30, 23], the modeling framework in these studies does not use the physical laws of volume conduction (Poisson’s equation) to predict measurements realistically, even though dipole approximation—weighted sums of the state variables of NMMs— have been used to model EEG[20].

The first objective of this study is to create a framework for modeling cortical column physics by embedding NMMs in a physical medium. Since synaptic and associated return currents in pyramidal cells are the main LFP generators [48, 16], we will assign spatial coordinates to apical and basal dendrites of the pyramidal populations corresponding to the locations where the flow of ions across the membrane takes place. Then, using Poisson’s equation (which governs the distribution of electrostatic potential in biological media) in a layered medium, we can realistically calculate the LFP profiles, bipolar LFPs, and CSD. We call this framework *laminar neural mass modeling*, or LaNMM for short, to emphasize its spatial and physical representation features.

As the first application of this approach, we explore a LaNMM adapted to simulate fast (gamma band) and slow (alpha band) dynamics, and we fit the model parameters to simulate multi-contact LFP recordings collected from the prefrontal cortex (PFC) of two macaque monkeys performing a working memory task. This previously collected dataset is described in Bastos et al. (2018) [6]. There, it was found that LFP power in the gamma band (30–250 Hz) was strongest at superficial layers and in the alpha/beta band (4–22 Hz) at deep layers and that the phase and amplitude from deep alpha drove the phase and amplitude of superficial gamma dynamics.

These findings align with other studies of the visual cortex of non-human primates [11, 15, 35, 57, 59, 28, 27]. However, the generality of these results has been recently questioned [18] since they may depend on the location of the recording site (e.g., visual vs. non-visual cortex), the task/stimuli type, and the type of measurement employed (e.g., CSD vs. LFP). Bollimunta et al. (2008) [12], using bipolar LFP and CSD measurements, found primary alpha power generators in the deep layers of the visual areas. However, in the inferior temporal gyrus (IT), alpha generators were located in superficial layers, and superficial to deep layer driving of alpha was found. Ninomyia et al. (2015) [47], also using bipolar LFP and CSD measurements, replicated the findings of Bastos et al. (2018) [6] in visual areas but not in the Supplementary Eye Field area. Finally, Haegens et al. (2015) [22] found maximal LFP alpha power in deep layers but a shift towards superficial layers using CSD. See Appendix A for a literature review summary of studies with different recording areas and measurement types.

A potential explanation for these discrepancies is that the LFP is calculated as the spatial line integral (along an arbitrary path) of the electric field from a (potentially remote) reference electrode to the recording site 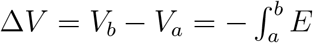, with the current density *J* = *σE*). Thus, LFPs are strongly susceptible to selecting the reference point and currents potentially far from the measurement point. This can affect the power distribution and coupling measurements (e.g., Granger causality). Using the recorded data, we study the impact of the choice of reference or measurement type on the electrophysiological power profiles. Ultimately, it would be desirable to avoid the ambiguity induced by LFPs by estimating more local quantities such as bipolar LFP—approximated as the first spatial derivative of the voltage along the linear array, which removes the referencing ambiguity but not volume conduction confounds—or the CSD—approximated as the second spatial derivative of the voltage multiplied by the tissue conductivity, which deals with both problems. CSD analysis reveals the location, direction (inwards or outwards), and strength of the flow of ions and is widely used to calibrate the laminar location of recording sites [36, 6, 19]. However, the derivatives (differences) computation can also decrease the signal-to-noise ratio.

Our second objective is to use our modeling framework to conduct a model-driven analysis of the data and spatially disentangle the slow and fast activity sources. By adjusting model parameters such as pyramidal synapse locations, we can adjust the LFP, bipolar LFP, and CSD power profile distributions and voltage correlations across the PFC laminae and compare them with the recordings collected in macaque monkeys by Bastos et al. (2018) [6]. To define a quantitative loss function for parameter fitting while avoiding referencing issues, we use the complete set of bipolar voltage correlations as the simulation target to maximize the correlation with the multi-contact data. With this loss function, we deduce a family of laminar models, composed of neuronal populations in superficial layers oscillating in the gamma band and neuronal populations in deep layers oscillating in the alpha band. The optimized architectures provide a mechanistic interpretation of the generation of slow and fast oscillations and approximate the measured LFP and CSD power profiles.

## 2. Methods

### 2.1. Multi-contact laminar recordings

The multi-contact dataset used in this study was collected in experiments described in Bastos et al. (2018) [6]. LFPs from the prefrontal cortex (ventrolateral prefrontal cortex and area 8a) of two macaque monkeys (*Macaca mulatta*) were recorded using a linear array of multi-contact laminar probes (16 contacts, 0.2 mm separation) while the animals were performing a search task (Figure 1A and 1B). All surgical and animal care procedures were approved by the Massachusetts Institute of Technology’s (MIT) Committee on Animal Care and were conducted following the guidelines of the National Institute of Health and MIT’s Department of Comparative Medicine. For our analysis, we selected the delay period (0.5—1s) of the successful memory encoding trials (see Appendix B for more details). The reference (ground) of the LFP recordings was located in the prefrontal cortex electrode chamber. To define which contacts belong to the superficial and deep layers, we aligned the electrode contacts where the cerebrospinal fluid (CSF) ends and the white matter (WM) begins. Our final selection included a total of 11 contacts spanning 2 mm of grey matter (GM), the first five contacts (0.8 mm) belonging to the superficial layers, and the rest in deep layers (1.2 mm), see Figure 1C and Figure 1E [6]. The data is not openly available but can be reasonably requested through a data sharing agreement to the corresponding author.

**Figure 1.**
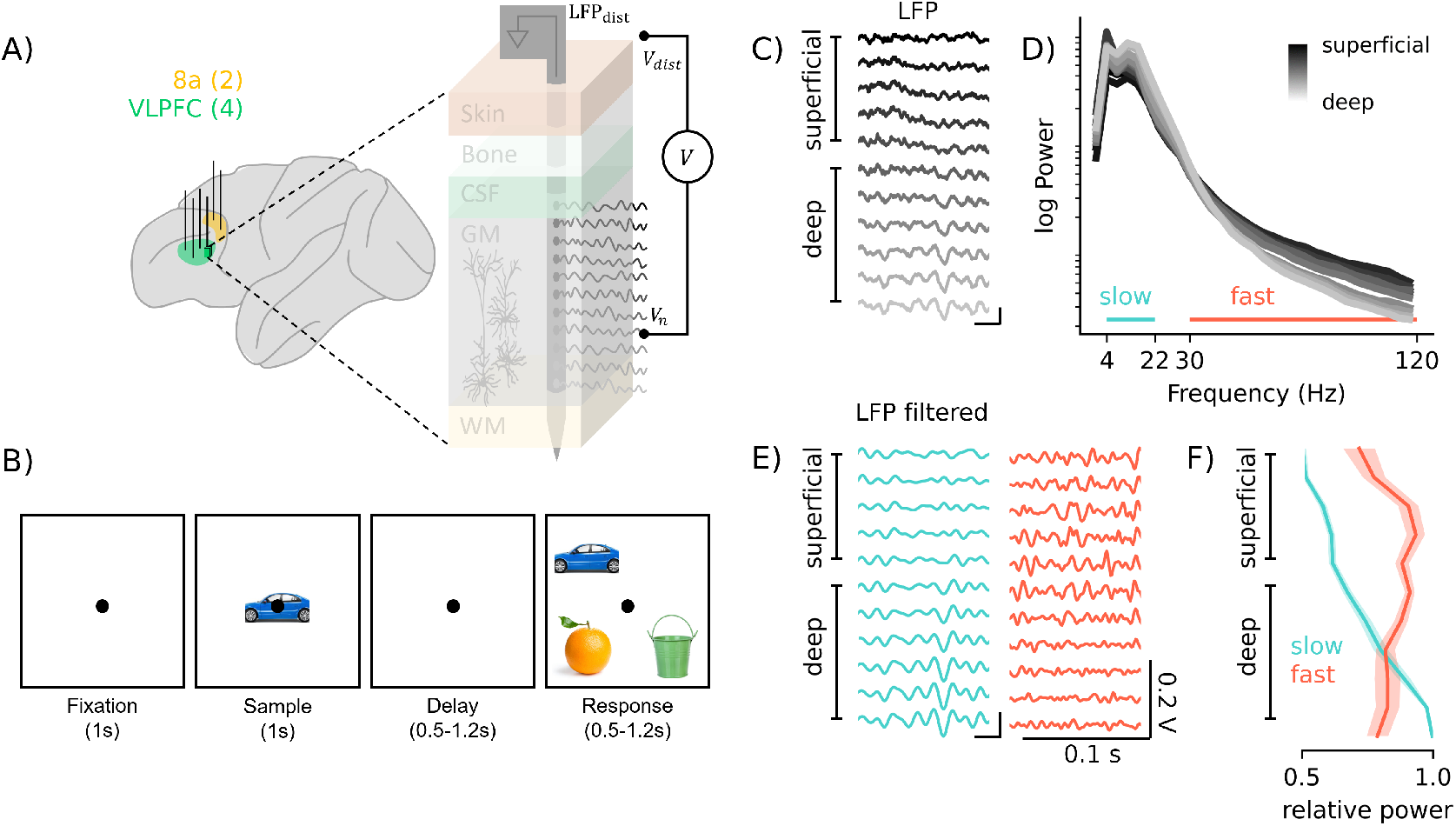
Multi-contact laminar recordings. A) Recordings from the ventrolateral prefrontal cortex (VLPFC) (4 electrodes) and area 8a (2 electrodes) referenced to the PFC electrode chamber. B) Visual search task. The macaque had to make a saccade to the match after the delay period (0.5–1.2 s). C) Sample LFP (referenced to ground in electrode chamber) recordings of the delay period for one of the trials. D) Sample power spectrum across depth (gradient from superficial-black to deep-white contacts). Sample in A) filtered in slow (4—22 Hz, blue) and fast (50—250 Hz) frequencies. Relative power across depth for the slow and fast frequencies.

Figure 1D displays the power spectral density across depth of the LFP data; there is a clear bump in slow frequencies (4–22 Hz), with a higher power in deep layers, and broadband gamma activity in fast frequencies (30–250 Hz, just until 120 Hz shown), with a higher power in superficial layers. We then filtered the data in these slow/fast frequency bands (Figure 1E) and computed the relative power across depth (Figure 1F). We found higher power in the fast band in superficial layers and the slow band in deep layers, as reported in Bastos et al. (2018) [6].

### 2.2. Synapse-driven formulation of NMM

Neural mass models (NMM) are mathematical representations of the dynamics of the average membrane potential and firing rate of a population of neurons in a cortical column [34]. In essence, a second-order differential equation describes the average membrane perturbation that a neuronal population *m* experiences at each synapse where it receives inputs from another population *n*. The synapse equation represents the conversion from an input presynaptic mean firing rate *φ*_*n*_ to a perturbation of the mean membrane potential *u*_*m←n*_ of the postsynaptic neuron population. We represent this relation here with the integral operator 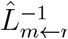 (a linear temporal filter), the inverse of which is a differential operator 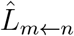,

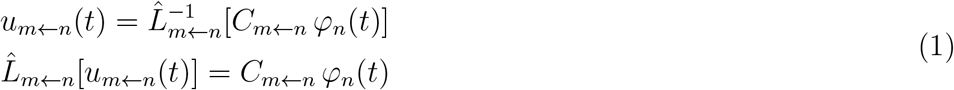

where *C*_*m←n*_ is the connectivity constant between the populations. The operator 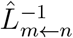 can be expressed as a convolution of the input signal with a kernel of the form *h*(*t*) = *Aat* exp[−*at*] for *t >* 0 [21], and satisfies the Green’s function equation 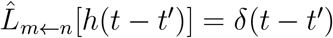 with appropriate causality boundary conditions.

For simplicity, we will use single index notation (*s*) to represent the synapse from one neuronal population to another, such that the set of the synapse transmembrane potential perturbations is {*s*} ≡ {*m* ← *n* : *C*_*m←n*_ ≠ 0}.

The linear operator that describes the dynamics of synapse *s* is defined as

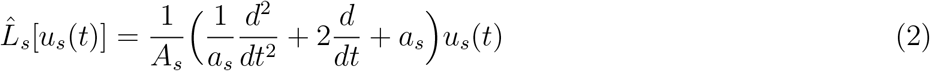

where *A*_*s*_ is the average excitatory/inhibitory synaptic gain and *a*_*s*_ is the rate constant of the synapse (*a*_*s*_ = 1*/τ*_*s*_, *τ*_*s*_ being the synaptic time constant).

Each neuronal population converts the sum *υ*_*m*_ of the membrane perturbations from each of the incoming synapses or external perturbations to an output firing rate (*φ*_*m*_) non-linearly by a sigmoid function,

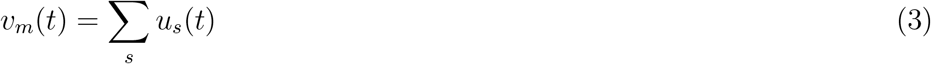

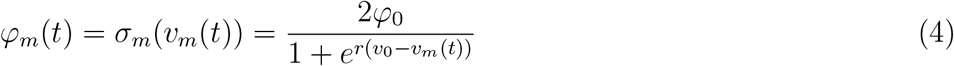

where *φ*_0_ is half of the maximum firing rate of each neuronal population, *υ*_0_ is the value of the potential when the firing rate is *φ*_0_ and *r* determines the slope of the sigmoid at the central symmetry point (*υ*_0_, *φ*_0_). We call this rewrite of the neural mass equations [24, 21] a synapse-driven formulation of an NMM. See Appendix C for more details.

### 2.3. Neural Mass Model

To generate the dynamics described in previous experimental studies [6, 35, 15, 57, 47, 13, 27], where the amplitude and phase of slow oscillations were observed to drive fast activity, we have combined two well-known NMMs. Slow oscillations in the alpha band (10 Hz) are produced by the Jansen and Rit model [24], and fast oscillations in the gamma band (40 Hz) by a variation of the PING model [14, 44].

The Jansen-Rit model (Figure 2) consists of a population *P*_1_ of pyramidal neurons, a population *SS* of excitatory cells (e.g., spiny stellate cells), and a population *SST* representing slow inhibitory interneurons (e.g., somatostatin-expressing cells, such as Martinotti cells). The PING model (Figure 2) consists of two populations: a pyramidal population *P*_2_, and a fast interneuron population *PV* (e.g., parvalbumin-positive cells, such as basket cells). The connectivity between these models is set so that there is a positive cross-frequency coupling and a negative power correlation from slow-to-fast frequencies, as is observed in experimental work [6]. Moreover, the connectivity profile is inspired by experimental and modeling studies characterizing fast and slow oscillations across the laminae ([38, 26] and the references therein).

**Figure 2.**
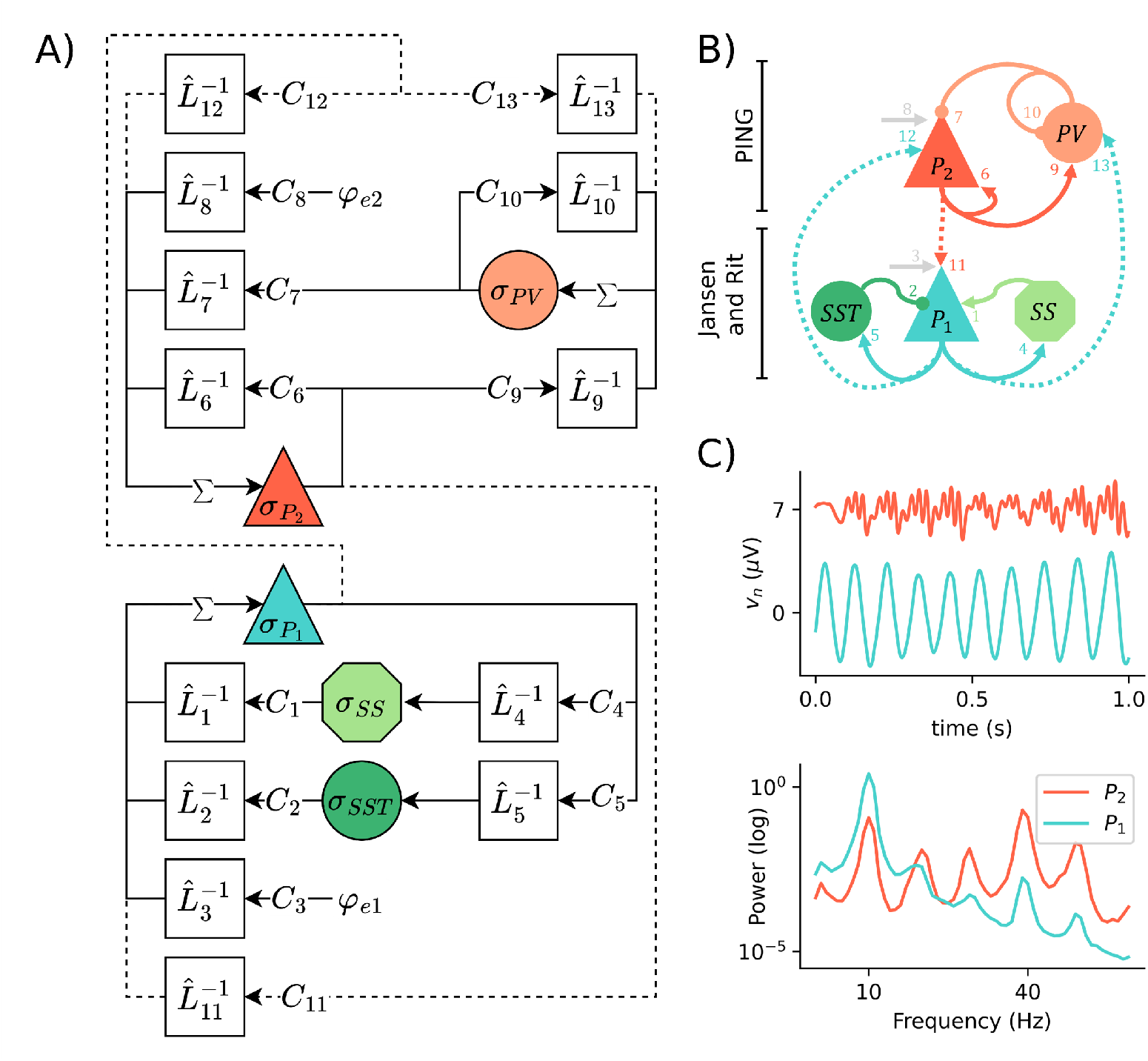
NMM for the macaque data. A) Diagram of the model equations. B) Illustration of the neuronal populations and the connectivity between them. Top, PING model, bottom, Jansen and Rit model. Rounded shapes in A) and B) represent inhibitory populations, the rest, excitatory. C) Top, membrane potential of the Pyramidal populations; bottom, power spectral density.

The model equations are visually represented in Figure 2A and described in detail, together with the parameters used, in Appendix D. Figure 2C shows the membrane potential and power spectra of the two pyramidal populations of the model.

### 2.4. Physical environment

In the laminar framework, we embed the NMM into a physical medium composed of two isotropic media—GM and CSF. We assume that the GM layers have a uniform thickness across depth, from 0 to 2 mm. To produce electrophysiological measurements from the model, we assume that synapses to pyramidal cells are the main current generators, given the anatomy of these cells (an elongated form factor), organization (perpendicular to the grey matter surface), and temporal coherence [48, 16].

The apical and basal dendrites of the pyramidal populations, with locations across the vertical *z*-axis (Figure 3A, *z*_*l*_ with layer *l* ∈ [1, 6]), provide the location of the input and output currents of each of the synapses (sinks and sources, respectively). For a detailed geometrical representation of the locations of the synapses with respect to the probe contacts across the GM see Appendix E. Since each synapse perturbation *u*_*s*_ has its location in space (*z*_*l*_), it will produce a flow of ions across the membrane, and therefore a synaptic current *I*_*s*_. We assume that the membrane perturbation of a given synapse *s, u*_*s*_, is linearly related to the injected current by a scaling factor that depends on the post-synaptic neuron population type and represents different aspects such as cell density and cell morphology. We capture this in a scaling factor—the *gain parameter g*_*n*_ —and write

**Figure 3.**
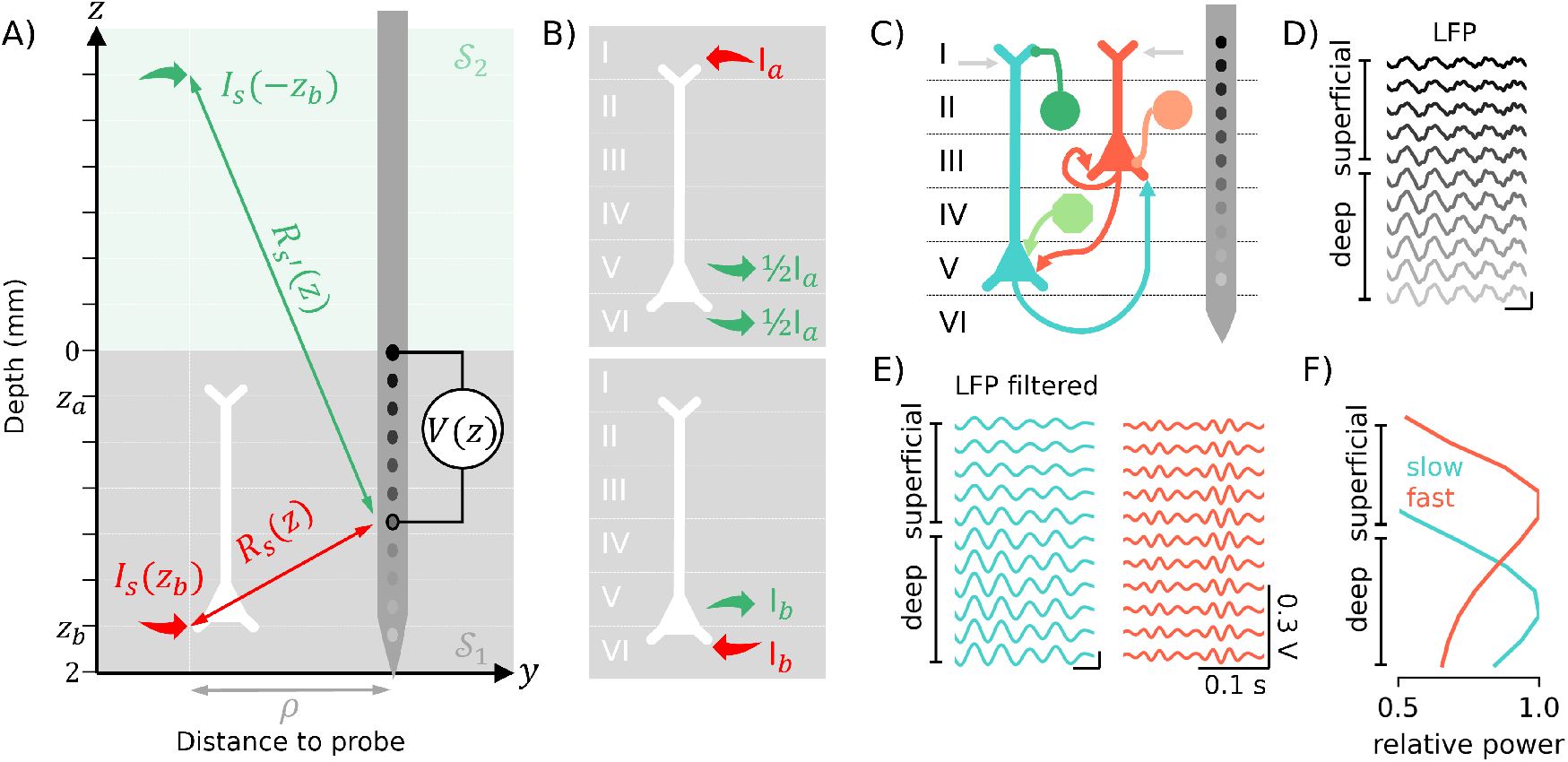
Laminar framework. A) Schematic of the geometry used to simulate LFPs. For simplicity, we have removed the time variable. B) Representation of the return currents for an apical synapse (top) and a basal synapse (bottom). C) Example of a biologically informed architecture (only the synapses to pyramidal populations are shown). D) LFPs generated by the architecture in C) for a probe distance *ρ* = 0.6 mm. E) LFP shown in D) filtered in slow and fast frequency bands. F) Relative power across depth for the LFPs shown in D) for the fast and slow frequency bands.

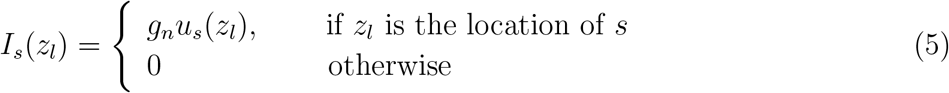

For simplicity, the time variable *t* is omitted here and in the following equations.

In the model, each injected current is accompanied by a capacitive return current (charge conservation) supplied by charges accumulated in the membrane. The precise flow of this current depends on cell morphology and electrical properties and is a subject of the study [39]. Based on previous studies [32], here we assume that inputs to apical dendrites (layer location *z*_*a*_) create a return CSD current at two locations on the basal dendrites, layer *z*_*b*_ and *z*_*b*+1_, each with half the total current since there are large dendritic ramifications at the soma of pyramidal neurons (Figure 3B). On the other hand, inputs to the basal dendrites (*z*_*b*_) create a return current at the layer above (*z*_*b*+1_). Thus, for each pyramidal cell, the total current generated at apical (ℐ_*a*_) and basal locations (ℐ_*b*_ and ℐ_*b*+1_) is

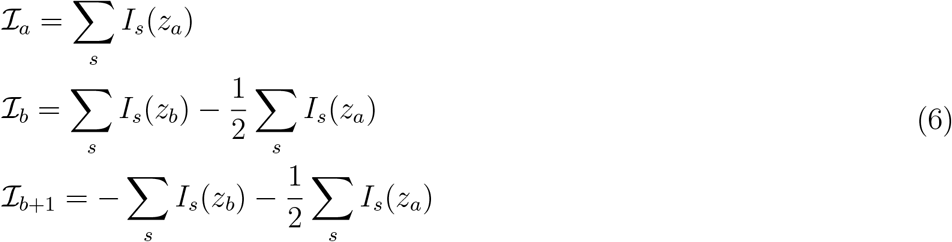

Once the current sources are specified, we can compute the electric potential and derived quantities. We model the potential field generated by each pyramidal cell by assuming there exist two isotropic media with conductivities *σ*_1_ = 0.40 S/m (GM) and *σ*_2_ = 1.79 S/m (CSF) [42] and a common planar boundary (Figure 3A). Then, the potential induced by a set of synaptic point current sources in GM is [48]

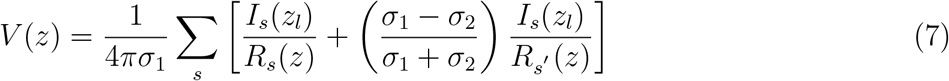

Here, *R*_*s*_(*z*) and *R*_*s*′_ (*z*) are the distances from the current source and mirror current source to the recording point (*z*), respectively (Figure 3A). These distances depend on the parameter *ρ*, representing the distance from the point source to the probe.

The normal component of the electric field can be computed from the gradient of the potential. We can simulate the experimentally measured “normal component” of CSD (A/m^3^) from the electric potential using

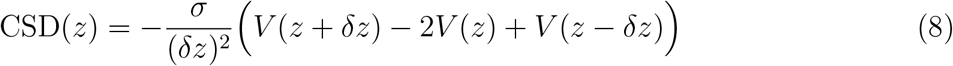

where *σ* (S/m) is the tissue conductivity [43, 50]. The values at the boundary layers are not evaluated.

### 2.5. Optimization function for model fitting

In order to compare the model and data and find optimal parameters, we computed the matrix of cross-contact correlations for the slow and fast frequency bands. The optimization process is represented in Figure 4 and described hereafter.

**Figure 4.**
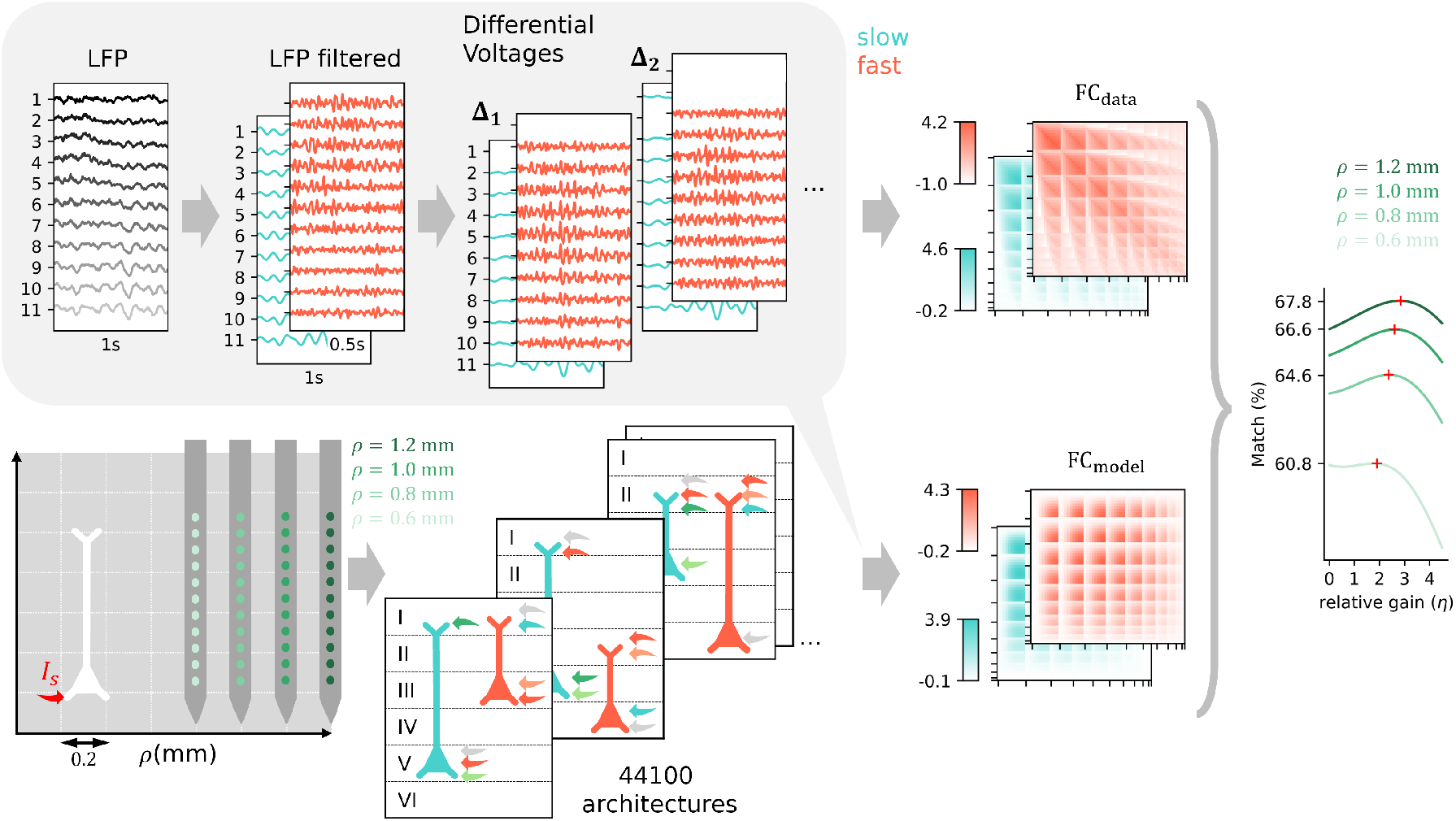
Overview of the optimization function for model fitting. The shaded box (top-left) shows the process from LFP data to the creation of the two-point function matrix, or functional connectivity matrix (FC, normalized by the standard deviation). To fit the model *FC* to the data, we optimized three different parameters: the distance to the probe *ρ*, the model architectures (a total of 44,100, just three samples represented), and the relative gain *η* (just one sample fit from all the architectures is shown here).

Let *V*_*a*_ be the empirical filtered measurement at a contact *a* referenced to ground. We then create the list of all bipolar combinations

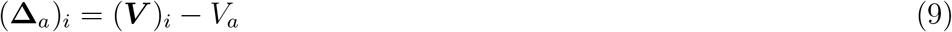

To avoid redundancy, in what follows, *i > a*. Then, to get a generalized reference-free functional connectivity representation (***FC***) matrix between all pairs of bipolar channels in the data, we compute the two-point function

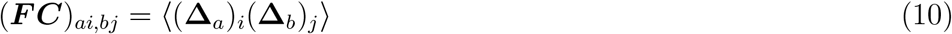

where the brackets denote the time average. Note that the two-point matrix ***FC*** includes as a subset the voltage power profiles referenced to any choice of the reference electrode.

As with real data, we can produce the two-point matrix from our model, ***FC***_*θ*_, that will depend on the parameters *θ*. We select the best model parameters by maximizing the Pearson correlation coefficient (*r*) between the flattened data ***FC***_*θ*_ and model ***FC*** matrices (only their diagonal and upper diagonal entries because they are symmetric) averaged over the two frequency bands, i.e., *θ*^*^ = arg max_*θ*_ χ(*θ*) with

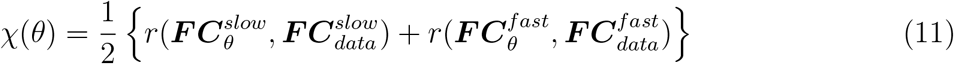

The results discussed below are provided as the percent match between the model and data (χ(*θ*) ∗ 100).

We adjusted the model using three sets of parameters (represented by *θ*): the distance to probe *ρ*, the model architecture, and the relative gain *η* between the slow and fast population. We explored 11 different *ρ* values, from 0.4 mm to 1.4 mm, with a size step of 0.1 mm (see Figure 4, for simplicity, just four values are shown). We also explored all possible model architectures by varying the location of every synapse to the pyramidal populations (a binary choice of either apical or basal location) and, thus, the location and layer span of the pyramidal populations. Figure 4 presents three examples of the different architectures. Each pyramidal population receives a total of 4 synapses to be assigned to one of six layers with the restriction that not all synapses in a population can be assigned to the same layer, so the analysis of combinations results in a total of ((6 · 5/2) (2^4^ − 2))^2^ = 44,100 possibilities. Here 6 · 5*/*2 is the number of possible apical/basal location pairs for a pyramidal cell population.At the same time, 2^4^ − 2 is the number of possible synapse assignments to each location, excluding the two cases where all the synapses are assigned to the same location (this ensures that a pyramidal population always spans two different locations). Finally, for each *ρ* and synaptic architecture choice, the relative gain factor between the slow and fast populations (*η* = *g*_*P*1_ */g*_*P*2_) was adjusted to maximize the optimization function. The *SciPy* library method *optimize* [58] was used to fit the model parameters.

## 3. Results

### 3.1. Optimization results

We first need to adjust model parameters to assess the model’s ability to simulate the power profiles for different LFP-derived measurements. The optimization function for model fitting is a generalization of the power profiles, namely the covariance of arbitrary differential voltage measurements (the two-point functions derived from bipolar voltages, see Methods section 2.5). We explored different model parameters to obtain the desired cross-correlation profiles.

The NMM parameters associated with intrinsic dynamics were fixed to produce representative fast (gamma) and slow (alpha) oscillations. We focused the fitting on laminar parameters directly influencing measurable quantities, namely the distance to probe (*ρ*), the location of the synapses (architectures), and the relative synaptic gain *η* (Figure 4 and F1). The laminar architecture parameters specify not only the location of the synapses in the pyramidal cells (apical vs. basal) but also the location of the apical and basal dendrites across the layers (I-VI).

The family of models that best fit the data is described in Figure 5. Figure 5A displays the percentage match with real data for different distances to probe *ρ* and the architectures ordered from worst to best match for an optimized relative gain (*η*). For most architectures, the best match happens with *ρ* = 1.4 mm, but if we zoom in to the 44 best architectures (0.1% of the total), we can see that the best fit happens with *ρ* = 1.0 mm. The model’s fit to the data degrades for *ρ >* 1.0 mm. It is noteworthy that the optimization function does not flatten out near the optimum, so the best solutions are sharply defined.

**Figure 5.**
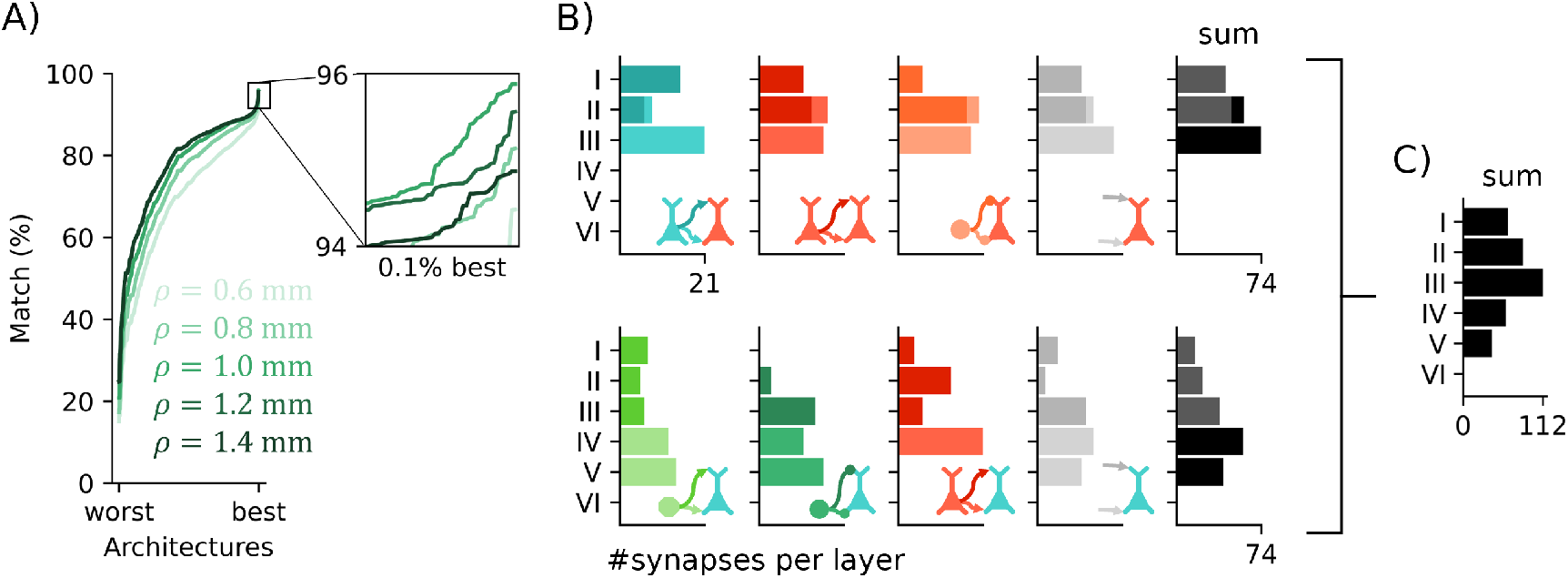
A) Percentage match with the real data for the different distances to probe *ρ* and ordered architectures from worst to best. B) Histograms of the number of synapses per layer among the best 0.1% architectures for each connection in the model with the best distance to probe (*ρ* = 1.0 mm). Synapses to the fast population are shown on the top row, and synapses to the slow population are shown on the bottom. Dark colors denote apical synapses and light colors basal synapses. The sum of the histograms for the synapse locations for both populations is shown for reference in black. C) Sum over all the synapses for both slow and fast populations.

Then, we analyzed the statistics of the resulting number of synapses per layer of the 0.1% best architectures (Figure 5B) for *ρ* = 1.0 mm, and the optimized relative gain (Figure F1). For the fast circuit, most of the synapses appear located in the superficial layers I–III, whereas the synapses of the slow circuit span layers I–V. The synapses to the basal dendrites for the slow population are located in layers IV and V, and in the fast circuit, the majority happens in layer III. In both the sum each connectivity plot, the peak synaptic activity for the slow population is always deeper than for the fast population. Although layer VI was included in the model, it was never the primary layer providing the synaptic currents in the top 0.1% of architectures. See Figure F1 for the optimized gains for the 0.1% best architectures.

### 3.2. Influence of the reference location in LFP measurements and model fit

To explore the effect of the electrical reference location on LFP measurements, we computed the LFP power profile using different electrical reference points: the ground in the prefrontal cortex chamber —a point distant to the sources (*LFP*_dist_— and the first superficial contact in the gray matter (*LFP*_0_). In order to mitigate the impact of possible far-field sources and referencing artifacts, we also evaluated the relative power of the fast and slow-frequency bands for the bipolar LFP and CSD measurements (Figure 6A).

**Figure 6.**
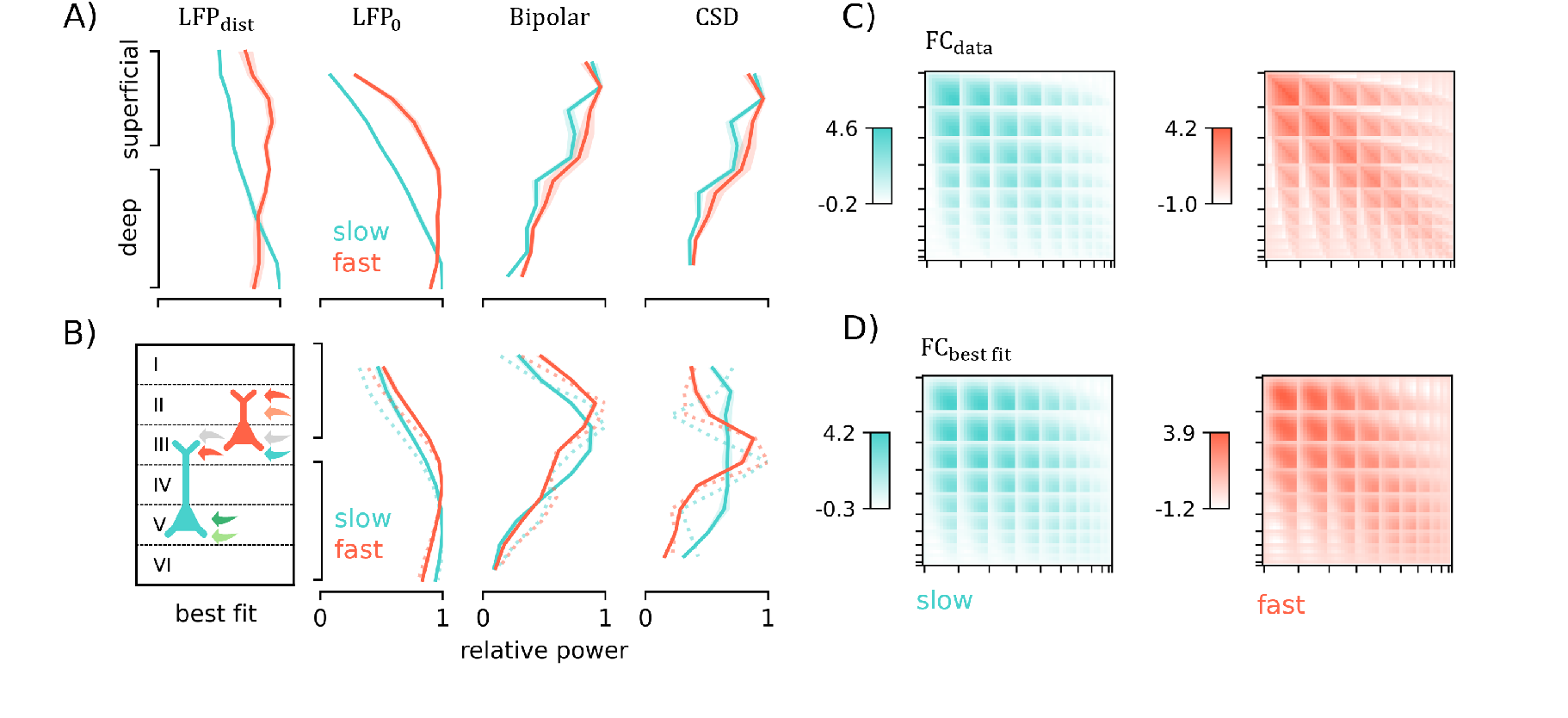
A) Relative power across depth for the real data. B) Relative power across depth for the average of the 0.1% best solutions in Figure 5. Each column shows the relative power profile for each measurement: LFP with a distant reference (*LPF*_dist_, not present in the model), LFP with the reference in the first contact (*LFP*_0_), bipolar LFP, and CSD. The dashed lines show the relative power of the best fit (architecture shown at the left). Filled areas show one standard error of the mean. Normalized FC by the standard deviation of C) the data and D) of the best model fit for the slow and fast frequency bands.

Given the definition of voltage as an integral of the field or currents and the results obtained from bipolar LFP and CSD measurements, we infer that the low-frequency LFP power data can be explained by currents generated by a long dipole spanning most of the cortex, with the high-frequency components generated by a shorter dipole in more superficial layers. Moreover, the *LFP*_0_ profiles in superficial layers display a rapid increase in power with depth compared with deep layers, where it slowly plateaus, which differs from the *LFP*_dist_ case, where the profiles remain more stable across layers for the fast frequency band, probably due to influences from more remote brain areas near the ground. This suggests that the spatial integral of the field along the vertical axis sums signals more coherently between the contacts in superficial layers than in deep layers, i.e., there is more spatial coherence of the electric field in superficial than deep layers. Moreover, the power peak of bipolar LFPs and CSD measurements also occurs in superficial layers.

We next computed the average relative power depth profile for the best 0.1% of architectures with *ρ* = 1.0 mm for each measurement (Figure 6B). These models also predict the rapid increase in power in superficial layers for *LPF*_0_ and the plateau in deep layers for both fast and slow frequencies. Furthermore, it shows a more superficial peak for the fast frequencies, as observed in the empirical data (Figure 6A, *LPF*_0_). The models also replicate the increased power in superficial layers for bipolar LFP and CSD. The estimated resulting density of synapses (Figure 5 C) is seen to reflect the associated CSD profiles from the model, with a peak in layer III (Figure 6 B, CSD).

The best solution is also shown in Figure 6B (best fit), in dashed lines, and together with the *FC*_*data*_ used for the model fitting (Figure 6C) as the best *FC*_*model*_ solution (Figure 6D). We observe that the model fits the slow-frequency *FC*_*data*_ better than the fast-frequency *FC*_*data*_. We also explored the power correlation and the modulation index of the model in Appendix G. We find similar patterns to those described in Bastos et al. (2018) [6], namely a bottom-up coupling of phase and amplitude and anti-correlation of fast and slow-frequency power.

Altogether, we conclude from fitting the data that our model’s main fast oscillatory synapses are located in superficial layers. This is consistent with the optimization results in Figure 5B, where most synapses were present in superficial layers in the fast frequency sub-circuit. In the slow population model fit, the synapses are located in significantly lower layers than the fast ones. Moreover, they span across almost all layers, peaking in layer IV with considerable synapse activity also in layer V, which is absent from the fast population.

## 4. Discussion

### 4.1. The relative power distribution across layers depends on the choice of the measurement

In addition to the importance of taking into account the reference point when studying data in LFP space, Figure 6A demonstrates that the distributions of the relative power vary significantly depending on the type of measurement. The CSD or bipolar LFP for the slow band peak shifts to superficial layers compared to voltage profiles, in agreement with other existing studies [11, 22]. We note that most studies that find the peak of slow oscillations in deep layers are based on monopolar LFP voltage recordings (*LFP*_dist_ here), with a remote reference far from the electrode contacts [6, 35, 15, 54, 28].

The referencing issue can, in part, be addressed by re-referencing the data to local electrode contacts [47, 28, 22]. It can be further mitigated using bipolar measurements (bipolar LFP, related to the local electric field and current density) or CSD estimates. Unlike CSD measurements, monopolar measurements (LFP) and bipolar LFPs are susceptible to volume conduction from remote sources since they are calculated as the spatial integral of the electric field between the measurement and reference point. It is critical to consider more local types of measurements, such as the bipolar LFP or CSD, to gain more information about the synaptic currents underlying LFP measurements [12, 22, 23] (see Figure A1).

It is important to note that other factors may influence the power profiles across layers: recording area [22, 47, 12], experimental task [28, 11, 22, 27, 18], and experimental procedures such as electrode placement [47]. The proper identification of the transition between superficial and deep layers, which should also depend on the area recorded [19, 15, 47], can also be confounding across studies. Future work should consider all these factors while trying to establish a golden standard for the experimental procedures (as suggested in [47]). The current availability of massively dense depth probe electrodes should shed light on these issues in the coming years.

### 4.2. Mesoscale laminar models can predict physical measurements of cortical rhythms across the laminae

A model-driven interpretation of the role of synaptic currents can shed some light on disagreements in the literature concerning the location of oscillatory generators. In addition to measurement issues related to referencing, confusion may arise from terminology—“generator” is a loose term that a physical modeling approach can clarify. Because electrophysiological recordings are driven synaptic currents which may be distant from the projecting or receiving cell bodies, there is a disassociation between soma location and generation locus. Thus, in this paper, we associate the term generators with synapses and the currents they generate.

In this study, we showed that our modeling framework could produce different oscillatory rhythms across layers and different types of laminar measurements extracted from multi-contact electrodes in a physically realistic manner (Figure 6). Despite the caveats listed in Section 4.1, it is interesting to note that we found the laminar generators for the slow rhythm to be located in significantly deeper layers than the generators of the fast rhythm. Indeed, the top-performing models for the fast (“gamma”) frequency synaptic generators included exclusively superficial layers 1–3. In contrast, the top performing models for the synaptic generators of the slower (“alpha”) frequency included deeper layers 4 and 5. This suggests that the generating circuit for alpha (and beta) oscillations integrates information across a larger spatial extent and samples from all layers.

### 4.3. Relation of the current model to current theories of alpha/beta and gamma oscillations

Our observation of superficial layers for gamma generation and superficial and deep layers for alpha/beta generation fits nicely with previous proposals of the role of these oscillations in the cortex. Alpha/beta has been implicated in feedback processes and gamma in feedforward sensory processing [7, 41]. Generally, top-down anatomical projections derive mostly from deep cortical layers, and bottom-up anatomical projections derive most strongly from superficial layers [37]. Bottom-up sensory processing is thought to rely on point-to-point connectivity and driving connections which determine the receptive field properties of downstream neurons [53]. Consistent with a bottom-up process, our model suggests that the gamma oscillatory circuit is largely constrained to the layers that send feedforward output (layers 2/3). Top-down processing is thought to rely on more modulatory, non-linear connections which integrate multiple streams of information [5]. Consistent with a more integrative, top-down process, our modeling results suggest that alpha/beta oscillations are generated by a more complex and spatially distributed process that may combine anatomical feedforward and feedback connections.

### 4.4. Limitations

Several limitations exist regarding the physical modeling framework we propose in this study. First, when estimating the voltage of the cortical column model (Equation 7), we assume the column can be represented as a set of monopoles and that the measurement point is relatively far from it. In reality, there is a field of dipoles in the cortical surface/patch, and the measurement contact can be placed precisely where the main dipole is located [16]. More realistic modeling approaches can be explored, such as describing the sources as homogeneous density distribution in the horizontal plane [39]. Additionally, as is often done in experimental work using depth probes, we assume that all currents occur in the vertical plane in our estimation of CSD from data. To properly extract the CSD, we need measurements in 3D space. However, this assumption is reasonable given that pyramidal cells are mostly homogeneously oriented perpendicular to the surface [16, 48].

Another limitation of this work is the relative simplicity of the presented NMM, with just two pyramidal populations oscillating in alpha and gamma bands, respectively. However, this simple model architecture inspired by the experimental work of Bastos et al. (2018) [6] has allowed us to explore all its combinations of pyramidal synapses. The LaNMM only allows for two synapse locations (apical and basal dendrites). Still, recent work that has addressed this limitation along with improving the model of return currents [39], which is also simplified in the present work.

## 5. Conclusions

In this study, we first extend the neural mass modeling formalism to include multiple oscillatory circuits and simulate realistic electrophysiological signals. We then use it to analyze data collected from multi-contact laminar measurements in the macaque. The analysis is performed with a simple laminar model whose connections are derived from literature and that is designed to produce coupled fast and slow oscillatory activity. We fit model parameters and the location of synapses by matching voltage statistics in the alpha/beta and gamma bands, leading to a solution with slow frequency oscillations generated by synapses spanning most cortical layers and fast oscillations in superficial layers. The laminar modeling framework developed here can help understand the neural mechanisms of electrophysiological signals and shed some light on controversial issues regarding discrepancies in LFP, bipolar LFP, and CSD measurements. The modeling framework may also help establish a firmer connection between neural mass models and EEG/MEG data and can be easily extended to analyze future data collected with dense probes. Finally, the possibility of modeling both slow and fast oscillatory activity within the same computational framework opens the possibility of understanding the origin of generalized EEG slowing observed in neurodegenerative conditions such as Alzheimer’s Disease and dementia, where slowing of alpha and reduced power of gamma activity are observed with disease onset and progression [9].

## Appendix A. Literature review

**Figure A1.**
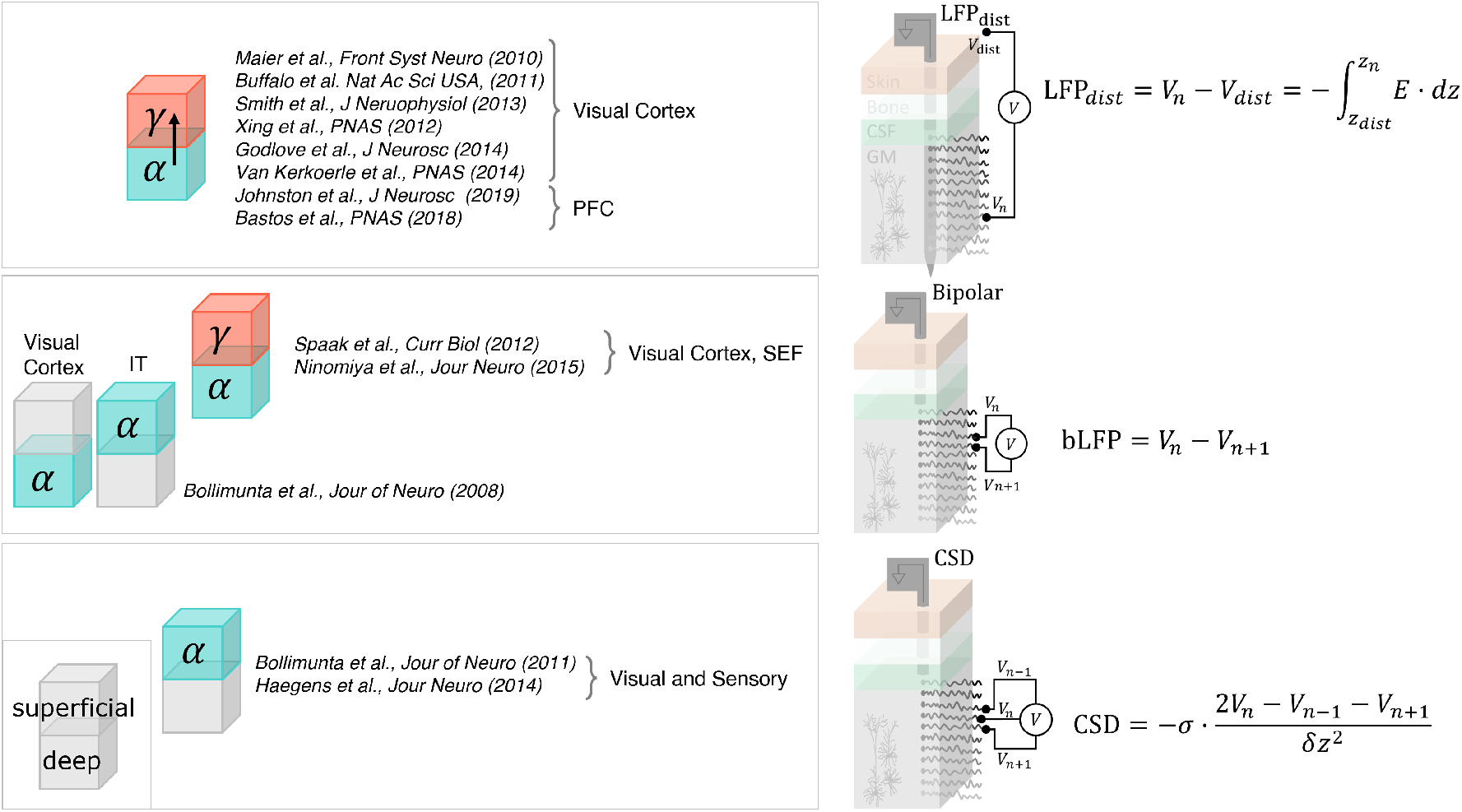
Summary of literature review of the slow (alpha/beta) and fast (gamma) cortical generators.

Figure A1 provides a graphical summary of the literature review performed of the different measurement types (rows) used in this study. It also shows the different results obtained for the different areas recorded. When using LFPs without taking into account reference location, most studies conclude that the fast activity is in upper layers and slow activity in lower layers. The conclusions change with other, more local measurement types (bipolar LFPs and CSD).

## Appendix B. LFP data analysis

The multi-contact dataset used in this study was collected in experiments described in Bastos et al. (2018) [6]. We analyzed the data from 2 monkeys (L, S, male and female, respectively) for six different sessions and two different brain areas (VLPFC and 8a). In Figure B1, the average across the correct trials for each session is shown. The delay period of monkey-L was fixed (1s), but for monkey-S it varied from 0.5 s to 1 s.

## Appendix C. Jansen and Rit model in synapse-driven formulation

### Appendix C.1. Jansen and Rit model description

In 1993, Jansen and Rit [25] developed a model of a cortical column which consists of three different neural populations: pyramidal neurons (*P*), inhibitory interneurons (*I*) and excitatory interneurons (*E*). The state variables of the model are the membrane potential and the firing rate of the neuron populations, and they are linked by two different transformations that shape the classical properties of neurons: the pulse-to-wave *h*(*t*) and wave-to-pulse *σ*(*υ*) functions [21, 2].

**Figure B1.**
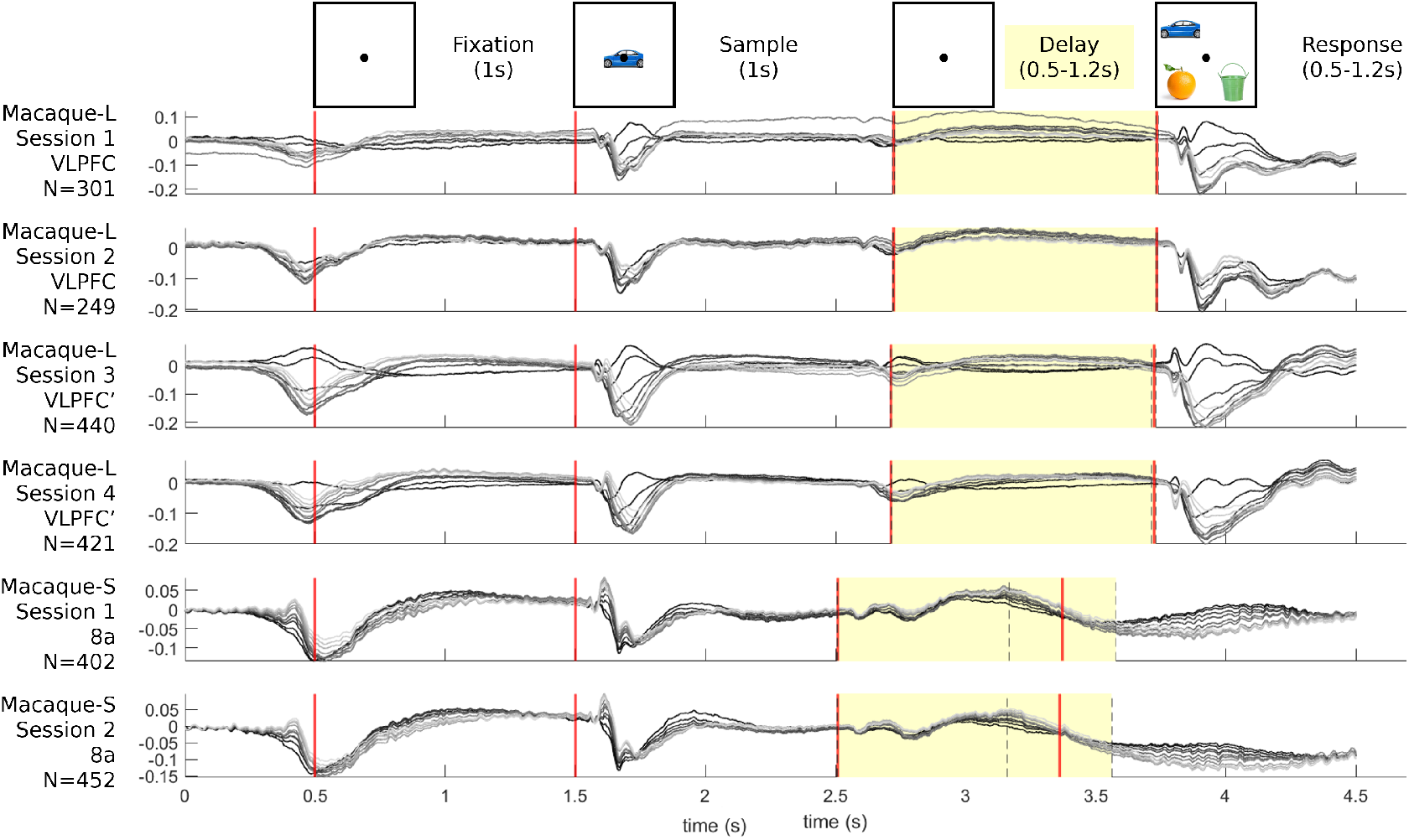
Average LPFs across trials, N, for the different monkeys and different sessions. The delay period is in between the ‘sampleOff’ and ‘testOn’ mark, in pink, which varies in time for monkey S (mean shown for the ‘testOn’, standard deviation in dashed lines). Each trace represent a different contact.

The *σ*(*υ*) operator, also called “wave-to-pulse”, introduces a nonlinear component that transforms the average membrane potential of a population *υ*(*t*) (m*υ*) into an average firing rate *φ*(*t*) (Hz)

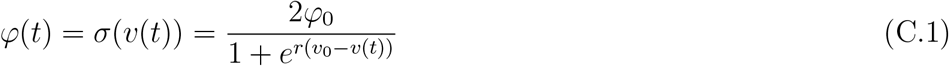

where *φ*_0_ is half of the maximum firing rate of each neuronal population, *υ*_0_ is the value of the potential when the firing rate is *φ*_0_ and *r* determines the slope of the sigmoid at the central symmetry point (*υ*_0_, *φ*_0_). See Table C1 for the standard parameter values of the model equations.

The *h*(*t*) operator, also called “pulse-to-wave”, converts the average rate of action potentials into an average post-synaptic potential, either excitatory *h*_0,1_(*t*) or inhibitory *h*_2_(*t*). The transformation is done by a second-order linear differential operator whose impulse response is given by

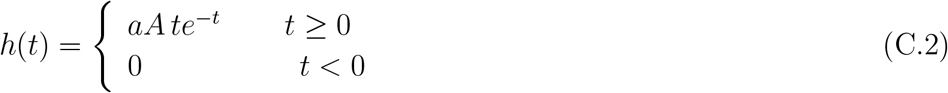

where *A* is the synaptic gain (in potential units, e.g., mV) and *a* (with units of time, s^*−*1^) is the rate constant (and its reciprocal *τ* the time constant) of the synapse. Each of these post-synaptic boxes corresponds to solving a differential equation of the form

**Table C1.** Parameters, description, and standard values of the JR synapse model. Values taken from [24]. Note that the sigmoid parameters in this model (*υ*_0_, *φ*_0_, *r*) are common to all neuron populations.

| Parameter | Description | Value |
| --- | --- | --- |
| $A_s$ | Average excitatory and inhibitory synaptic gain | $A_{1,3,4,5} = 3.25 \text{ mV}$<br>$A_2 = -22 \text{ mV}$ |
| $a_s$ | Time rate constant of average excitatory and inhibitory postsynaptic potentials | $a_{1,3,4,5} = 100 \text{ s}^{-1}$<br>$a_2 = 50 \text{ s}^{-1}$ |
| $C_s$ | Average number of synaptic contacts between population types | $C_1 = 108$<br>$C_2 = 33.7$<br>$C_3 = 1$<br>$C_4 = 135$<br>$C_5 = 33.75$ |
| $v_0$ | Potential when 50% of the firing rate is achieved | 6 mV |
| $\varphi_0$ | Half of the maximum firing rate | 2.5 Hz |
| $r$ | Slope of the sigmoid function at $v_0$ | $0.56 \text{ mV}^{-1}$ |

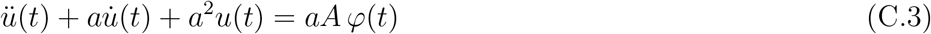

where *φ*(*t*) is the output of the sigmoid function (average firing rate of a population, in Hz) and *u*(*t*) is the membrane potential alteration in each of the synapses. The function *h*(*t*) is the equation’s Green’s function or impulse response, i.e., the solution with *φ* = *δ*(*t*) and appropriate boundary conditions. The parameters *A* and *τ* = 1*/a* represent the maximal amplitude of excitatory or inhibitory post-synaptic potential and the average time constant for each synapse type, respectively.

This second-order differential equation can be decomposed in a system of two equations,

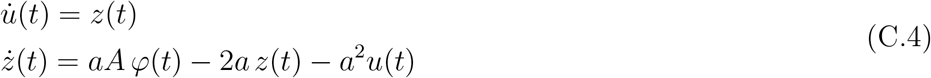

There are thus three main state variables in the model: the average membrane potential of each of the subpopulations of the system: *υ*_*P*_ (*t*) for the pyramidal cells, and *υ*_*E*_(*t*), *υ*_*I*_(*t*) for the excitatory and inhibitory interneurons, respectively. The average membrane potential *υ*_*P*_ of the pyramidal population has been typically used as a proxy source of electrophysiological signals such as LFPs and EEG (dipole generator). We improved upon this first-order approximation in the LaNMM framework (see next section).

The Jansen and Rit model can be described with a set of six differential equations with each pair corresponding to a population,

**Figure C1.**
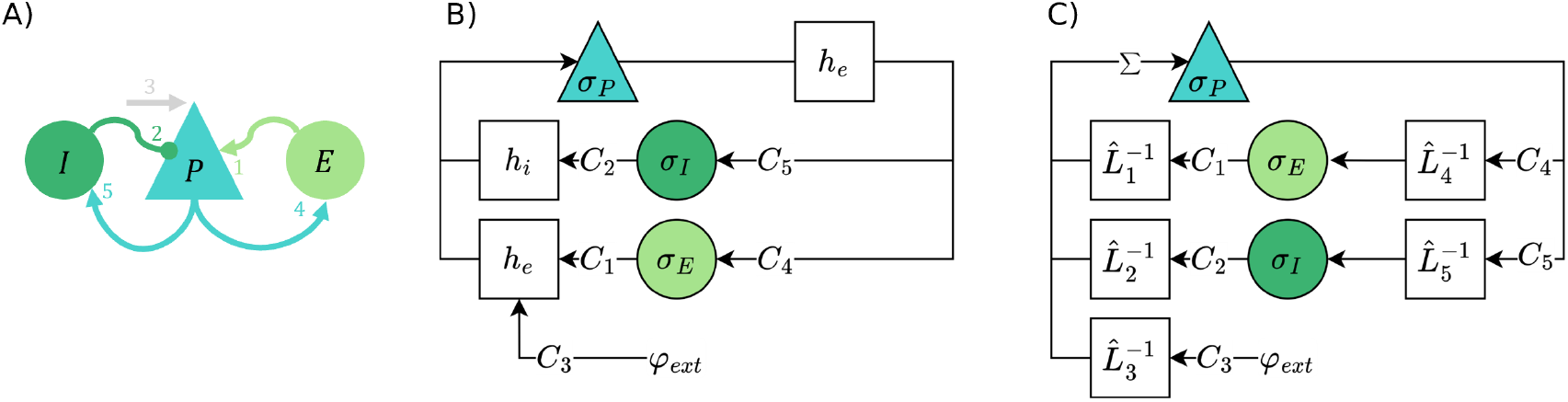
Jansen and Rit model. A) Schematics of the connections between the different populations: P—Pyramidal, I—Inhibitory interneuron and E—Excitatory interneuron. B) simplified wired diagram exploiting the presence of common synapse types, c) full synapse-driven wiring diagram with all synapses explicitly represented.

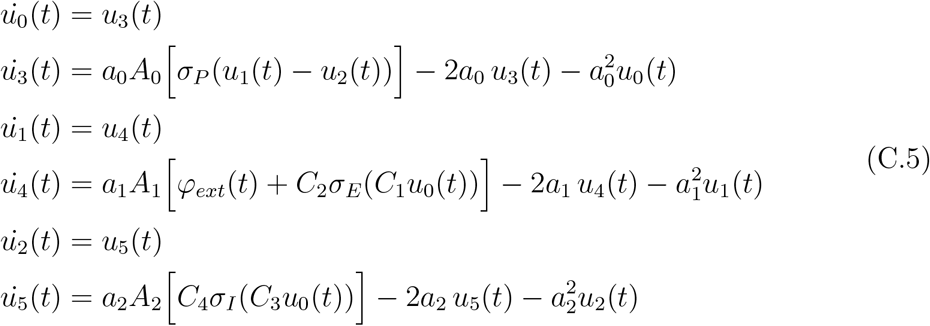

For an illustrative description of the model equations see Figure C1 A) and B). In this cortical column configuration, the membrane potential of the pyramidal population is *υ*_*P*_ (*t*) = *u*_1_(*t*)−*u*_2_(*t*), the membrane potential of the inhibitory interneuron population is *υ*_*I*_ = *u*_0_ and of the excitatory interneuron population is *υ*_*E*_ = *u*_0_.

### Appendix C.2. Derivation of the synapse-driven formulation from Jansen and Rit equations

We can rewrite the Jansen and Rit NMM focusing on the dynamics of each of the synapses independently, which will allow us to generalize the equations and simplify the definition of the neural dynamics for the development of more complex models. We will define a new linear operator, 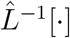[·], to transform the pre-synaptic average firing rate of neuron *n φ*_*n*_ into a post-synaptic membrane perturbation of neuron *m u*_*m←n*_:

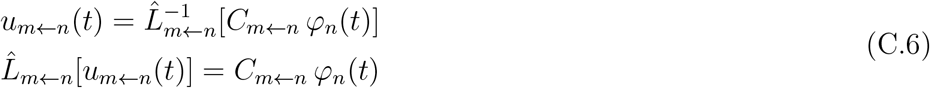

The inverse of the 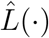 operator and can be expressed as an integral (convolution) operator using the typical *h*(*t*) kernel,

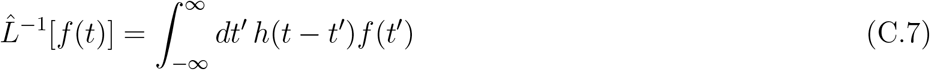

Note that, for simplicity, the index *s* will represent the synapse from one neuronal population to another *m* ← *n*, where *n, m* ∈ [*P, E, I, ext*] and (*m, n*): *C*_*m←n*_ ≠ 0. Then, we can define the linear operator that captures the synapse dynamics 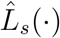 as

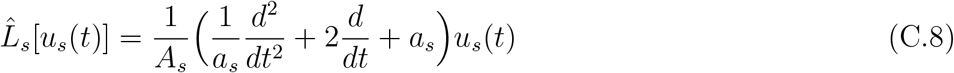

The sum of each pre-synaptic perturbation into neuron *n* is the overall membrane potential perturbation of the post-synaptic neuron, *υ*_*m*_,

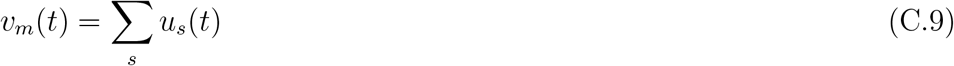

and the average firing rate of the neural population, *φ*_*m*_, is the output of the non-linear function,

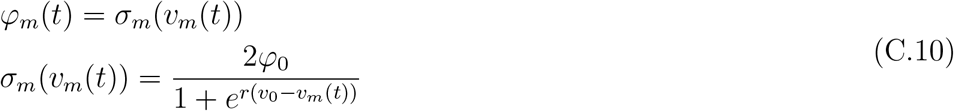

Finally, the set of equations, one for each synapse (*m, n*) and neuron *m*,

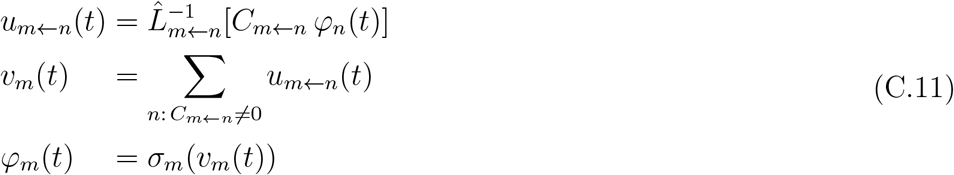

which we call the synapse-driven reformulation of the Jansen-Rit that can be easily be extended to other, more complex NMMs.

Rewritten using the synapse-driven formalism, the Jansen and Rit equations specify the dynamics as a function of the average firing rate for each neural population *φ*_*n*_, the average membrane potential for each population *υ*_*n*_, and the membrane perturbation per each synapse *u*_*s*_,

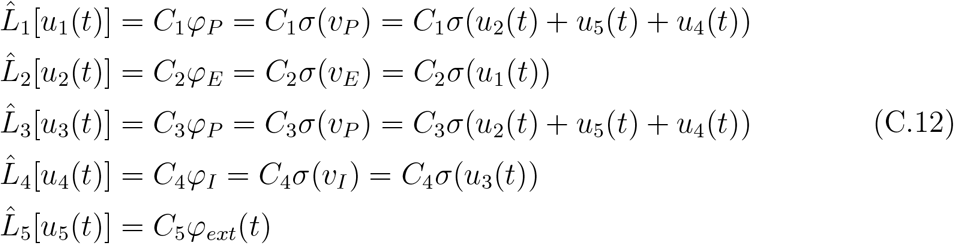

Figure C1 C) provides the diagram and dynamics of the Jansen and Rit NMM in the Synapse-driven implementation.

## Appendix D. Model parameters and equations

The parameters of the model are described in Table D1 and the equations are the following:

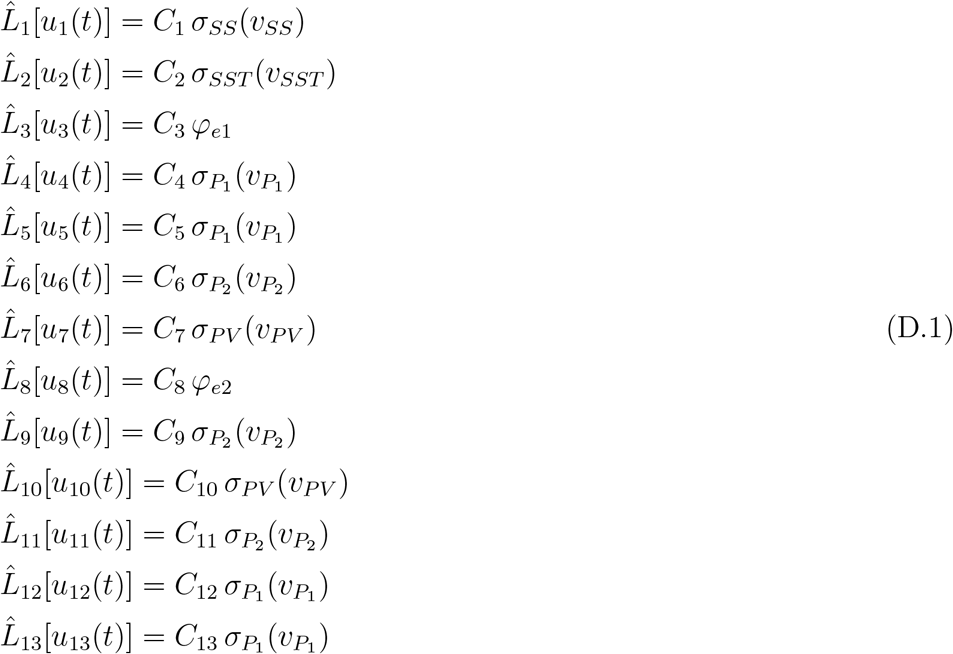

with neuronal population membrane potentials given by:

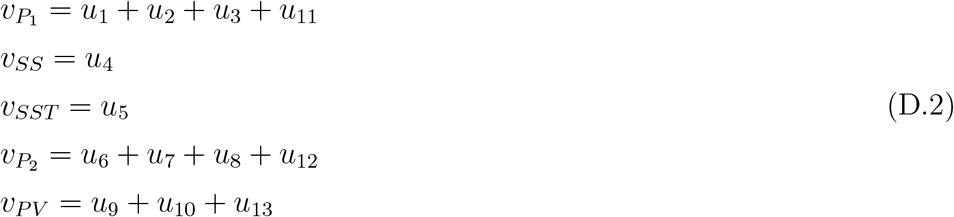

## Appendix E. Geometrical representation of the laminar model

In this section we show the location of the synapses used in the LaNMM together with the contact location of the different electrode positions in the GM (Figure E1). The cortical depth (2 mm) is shown in the shaded area, where all the sources and contacts are located. The sources (N=6) and contacts (N=11) are equally distributed in the cortical space. We also show the different distances to probes *ρ* used in the model fitting.

## Appendix F. LaNMM contact couplings

The median relative gain of the 0.1% best architectures for *ρ* = 1.0 mm is 0.13, meaning that the gain of the fast circuit *g*_*P*2_ is approximately seven times higher than the gain of the slow circuit *g*_*P*1_, *η* = *g*_*P*1_ */g*_*P*2_. This might be due to the fact that the intrinsic power of gamma oscillations in our model is one order of magnitude lower than the power of alpha oscillations (Figure F1).

**Table D1.** Parameters, description, and standard values of the model. Values are taken from [24] and [44]. Note that the sigmoid parameters in this model (*υ*_0_, *φ*_0_, *r*) are common to all neuron populations. Moreover, excitatory synapses have the same synapse dynamics, (*A, a*)_*AMP A*_ = (*A, a*)_1,3,4,5,6,8,9,11,12,13_, and inhibitory synapses have either fast dynamics, (*A, a*)_*GABAfast*_ = (*A, a*)_7,10_, or slow, (*A, a*)_*GABAslow*_ = (*A, a*)_2_.

| Parameter | Description | Value |
| --- | --- | --- |
| $A_s$ | Average excitatory and inhibitory synaptic gain | $A_{AMPA} = 3.25$ mV<br>$A_{GABA_{slow}} = -22$ mV<br>$A_{GABA_{fast}} = -30$ mV |
| $a_s$ | Time rate constant of average excitatory and inhibitory postsynaptic potentials | $a_{AMPA} = 100$ s<sup>-1</sup><br>$a_{GABA_{slow}} = 50$ s<sup>-1</sup><br>$a_{GABA_{fast}} = 220$ s<sup>-1</sup> |
| $C_s$ | Average number of synaptic contacts between population types | $C_1 = 108, C_2 = 33.7$<br>$C_3 = 1, C_4 = 135$<br>$C_5 = 33.75, C_6 = 70$<br>$C_7 = 550, C_8 = 1$<br>$C_9 = 200, C_{10} = 100$<br>$C_{11} = 80, C_{12} = 200$<br>$C_{13} = 30$ |
| $v_0$ | Potential when 50% of the firing rate is achieved | 6 mV<br>except for $P_2$ : 1mV |
| $\varphi_0$ | Half of the maximum firing rate | 2.5 Hz |
| $r$ | Slope of the sigmoid function at $v_0$ | 0.56 mV<sup>-1</sup> |
| $\varphi_e$ | External input | $\varphi_{e1}$: pink noise with mean 200 Hz and standard deviation 30 Hz<br>$\varphi_{e2} = 90$ Hz |

## Appendix G. LaNMM contact couplings

Here we show the power correlation and the modulation index (MI) for the model LFP data. The power correlation is computed by extracting the amplitude of the band-passed signals using the Hilbert transform and then computing the Spearman correlation of the envelopes. The MI is computed as the entropy of the phase-amplitude histogram, with phase measured in the slow band and amplitude in the gamma band.

We show in Figure G1 that there is a generic negative power correlation between contacts. The most negative peak happens between the deep slow band and the superficial fast band for the best fit model parameters (box 4), and for the average over the best 0.1% model parameters from superficial to superficial layers (box 1).

**Figure E1.**
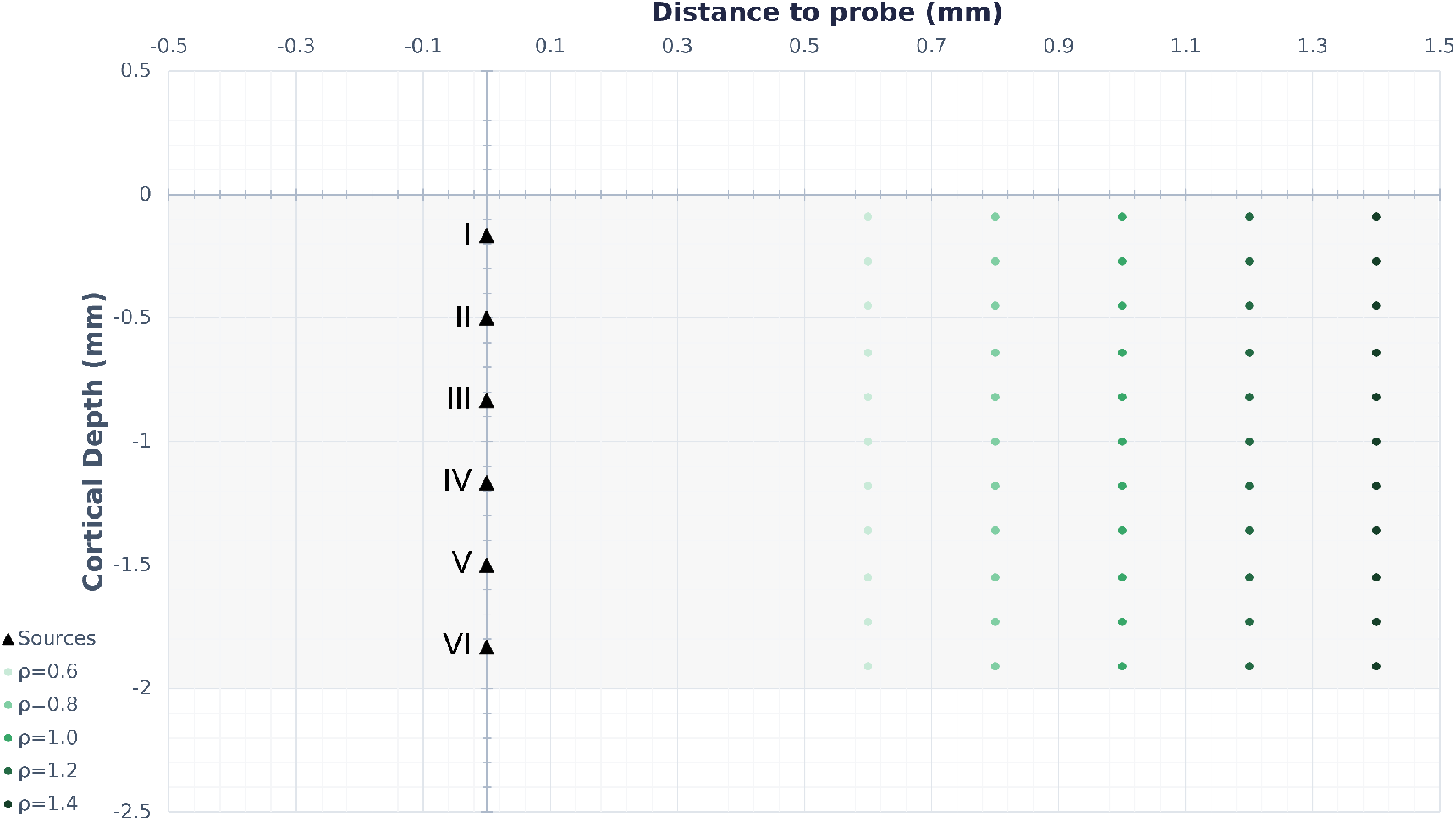
Schematic of the locations of the sources in every layer and the probe contacts for the different distances to probe (*ρ*) in the grey matter (2 mm).

**Figure F1.**
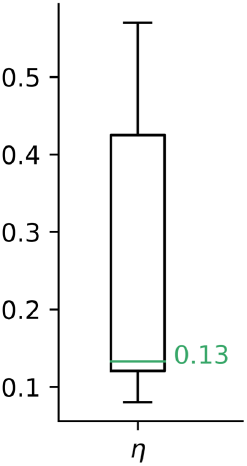
Box plot of the optimized relative gain *η* with the median of the distribution in green for the best 0.1% architectures in Figure 5.

In the case of MI, the model always displays a positive MI through all the contacts, with the peak in box 1, from superficial slow frequencies to superficial fast ones.

**Figure G1.**
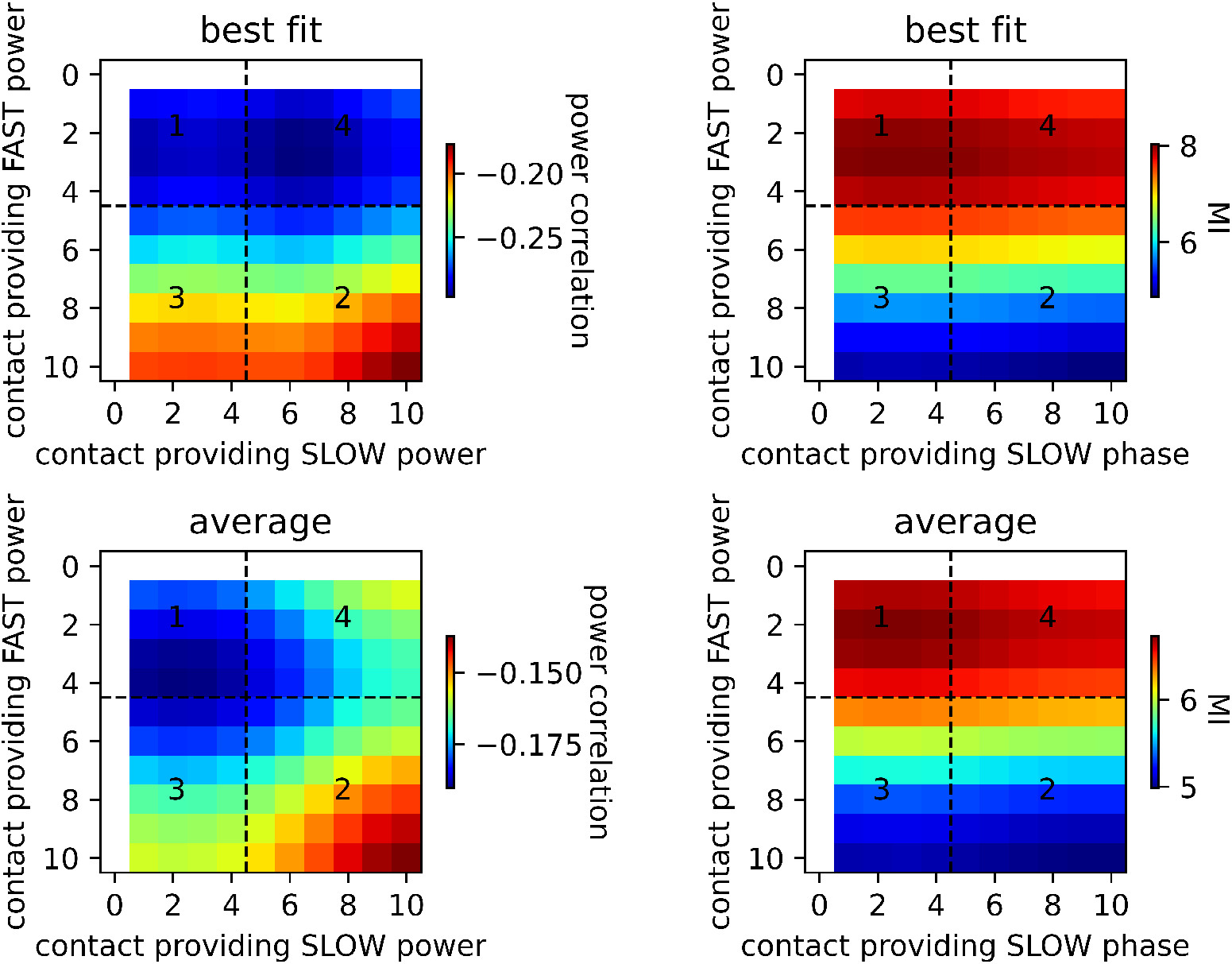
Power correlation (left) and modulation index (right) for the best model fit (top) and averaged over the best 0.1% model results. The dashed lines refer to the transitions between superficial and deep layers. The white rows indicate the reference contact.

## Author contributions

**Roser Sanchez-Todo**: Conceptualization, Methodology, Software, Validation, Data Curation, Writing-Original Draft, Visualization; **André M. Bastos**: Conceptualization, Resources, Data Curation, Writing-Review and Editing; **Edmundo Lopez Sola**: Software, Writing-Review and Editing; **Borja Mercadal**: Conceptualization, Methodology; **Emiliano Santarnecchi**: Writing-Review and Editing; **Earl K. Miller**: Writing-Review and Editing; **Gustavo Deco**: Supervision, Writing-Review and Editing; **Giulio Ruffini**: Conceptualization, Methodology, Formal analysis, Writing-Original Draft, Supervision, Project administration.

## Acknowledgments

This work has received funding from FET European Union’s Horizon 2020 research and innovation programme (grant agreement No 101017716). The authors wish to thank Esteban Félez and Kyra Kadhim for revising the document, and Fabrice Wendling for useful discussions.

## Notes

### Competing Interest Statement

RST, GR, BM, ELS work for Neuroelectrics, a company dedicated to the creation of brain stimulation solutions. GR is a cofounder of Neuroelectrics.

### Summary of Updates

Manuscript polishing, added highlights.

